# Establishing defined daily doses (DDDs) for antimicrobial agents used for pigs, cattle and poultry in Japan and comparing them with European DDD values

**DOI:** 10.1101/2020.12.23.424121

**Authors:** Kyoko Fujimoto, Mai Kawasaki, Reiko Abe, Takashi Yokoyama, Takeshi Haga, Katsuaki Sugiura

## Abstract

Monitoring of antimicrobial use is essential to manage the development and selection of antimicrobial resistance. A variety of indicators has become available to monitor antimicrobial use in human and animal medicine. One of them is an indicator based on defined daily dose (DDD). By using the number of DDDs used and normalizing it by the population at risk of being treated in a defined period, one can estimate the number of treatment days with antimicrobial agents in a population. For veterinary medicine, the European Medicines Agency (EMA) has published the European values of DDD (DDDvet) for food-producing animals. In this study, we defined Japanese defined daily doses for antimicrobial agents (DDDjp) using DDD values that we previously assigned for antimicrobial products approved for use in pigs, cattle and poultry in Japan and compared them with DDDvet values. For the comparison, the quotient of Japanese and European values (QDDD) was calculated and the effect of the administration route and the number of active substances contained in the preparation was investigated. A total of 59, 51 and 27 DDDjp values were defined for 43, 32 and 25 antimicrobial agents using the data of 269, 195 and 131 products approved for use in pigs, cattle and poultry respectively. A comparison was possible for 44, 27 and 17 pairs of DDDjp and DDDvet values for antimicrobial agents used for pigs, cattle and poultry respectively. The comparison showed median QDDD value of 0.66 and 0.63 for antimicrobial agents used for pigs and cattle respectively (P<0.01), indicating that the Japanese daily doses are significantly lower than the corresponding EMA values in these species. For the antimicrobial agents used for poultry, no siginificant difference was observed between DDDjp and DDDvet values with median QDDD value of 1.17. The difference between DDDvet and DDDjp values and absence of DDDvet values for some antimicrobial agents marketed in Japan indicate that DDDjp rather than DDDvet should be used as the basis for the calculation of antimicrobial use monitoring in farm animals in Japan.

## Introduction

The use of antimicrobial agents in food producing animals may lead to the emergence and selection of resistant bacteria. The bacterial resistance arises through complex mechanisms, normally through mutation and selection, or by acquiring from other bacteria the genetic information that encodes resistance [1]. Therefore, reducing the selection pressure by reducing antimicrobial usage is considered one of the important strategies to decrease the resistance rate [1].

An important tool for the control and reduction of antimicrobial use in veterinary medicine is the establishment of a monitoring system, which can be based on various types of data collection [2]. In the EU, antimicrobial sales data are collected from each member country and antimicrobial consumption in each member is calculated and published in terms of milligrams of active ingredient sold per population correction unit (mg/PCU). The authors have previously calculated the antimicrobial usage in food-producing animals in Japan using the same indicator [3]. The disadvantage of this monitoring system is that the different potencies of different antimicrobial agents are not taken into account [4]. In human medicine the World Health Organization (WHO) has determined an average daily maintenance dose for the main indication for each active substance [5]. Using the daily doses and the amount of an active ingredient used, the number of potential treatment days in a population can be estimated. This statistical value has been adapted to veterinary medicine and is the basis of the national antibiotic monitoring systems in several Scandinavian countries and the Netherlands [6, 7, 8].

A list of defined daily doses for the food-producing animals (DDDvet) has been available from the European Medicines Agency (EMA) since 2016 [9]. Dose data from nine EU member states were collected and average values for the daily doses of each active ingredient by administration route (“parenteral”, “oral except premix” and “premix”) and by species were calculated. The antimicrobial active ingredients were classified based on anatomical-therapeutic-chemical correspondences (ATCvet Code) [10].

In the hope to establish a monitoring system using an indicator based on daily dosage, the authors have previously assigned DDD values for 269, 195 and 131 veterinary antimicrobial products approved and marketed for use in pigs, cattle and poultry in Japan [11, 12].

The aim of the present study was to define Japanese daily doses (DDDjp) for each antimicrobial agent (active ingredient) based on these DDD values assigned for products and to compare them with the values of the EMA.

## Materials and Methods

### Defining Japanese DDD values for antimicrobial agents used for pigs, cattle and poultry in Japan (DDDjp)

The DDDjp values were calculated using the DDD values that we previously assigned for 259, 195 and 131 veterinary antimicrobial products approved and marketed for use in pigs, cattle and poultry respectively in Japan [11, 12]. In our previous study, the DDD values for antimicrobial products other than intramammary and intrauterine products were assigned by species and by kg animal, using the principles developed by the European Medicines Agency [13]. The DDD values were assigned by kg animal by dividing the daily dose by 635kg (average weight of dairy cow in Japan) for intramammary products for lactating cows and intrauterine products, and by multiplying the dose by 4 (number of teats) and dividing it by 635kg and an assumed long acting factor of 10 days for products for dry cows [12].

In the current study, the DDDjp values were calculated by averaging the DDD values of products if there are two or more products containing the same antimicrobial agent:

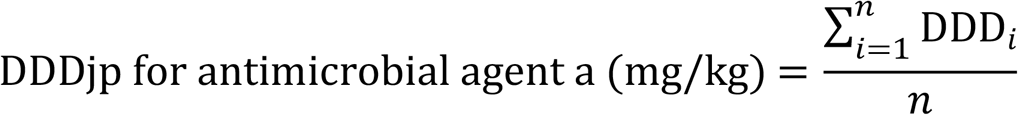

where DDDi is the DDD value (mg/kg) of antimicrobial product i containing antimicrobial agent a, and n is the number of products containing antimicrobial agent a. For those antimicrobial agents that are used as active ingredient in products for two or more administration routes, DDD values were assigned separately by administration route. Likewise, for those that are used both in single substance and combination products, DDD values were assigned separately by the number of active ingredients contained in the preparation. In other words, the average (arithmetic mean) of all DDD values of products for each combination of species, antimicrobial agent, administration route and the number of substances in the product (single substance or combination product) was used to assign DDDjp – e.g. pig/benzylpenicillin/injectable/single substance.

### Comparison of DDDjp and DDDvet values

For the comparison with the values of the EMA, the quotient of the daily doses (QDDD) was formed from Japanese and EMA values:

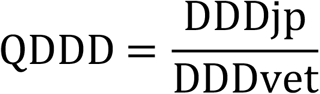

If the quotient shows a value of one, it means that the Japanese dose and the EMA dose are the same. Quotients greater than one mean that the Japanese dosages are higher and values below one mean that they are lower. If there was no EMA value for an active ingredient, the antimicrobial agent was excluded from the comparison. The effect of the number of active ingredients contained in the product and the administration route on the comparison was examined.

### Statistical analysis

The statistical analysis was carried out to verify a deviation of QDDD between administration routes and number of active ingredients with BellCurve for Excel version 3.20 (Social Survey Research Information Co., Ltd.). A P-value ≤ 0.05 was set as the significance level. The data were checked for normal distribution by Shapiro tests. The difference between DDDvet and DDDjp values was examined using the Wilcoxon test for paired samples. The effects of the administration routes and the number of active ingredients in the product were examined using a Mann-Whitney U test.

## RESULTS

### Distribution of antimicrobial products approved for use in pigs in Japan

A total of 60, 54 and 27 DDDjp values were defined for 43, 32 and 25 antimicrobial agents using the data of 259, 195 and 131 products approved for use in pigs, cattle and poultry respectively. The distribution of the products according to the administration route and the number of active substances in the preparation is shown in Tables 1, 2 and 3. A complete list of the antimicrobials for which DDDjp values were defined are shown in Table 4.

**Table 1.** Distribution of the antimicrobial products approved for use in pigs in Japan by administration route and the number of active ingredient contained

| Type of product | Administration route |  |  | Total |
| --- | --- | --- | --- | --- |
|  | Injection | Oral | Topical |  |
| Single substance product | 75 | 133 | 1 | 209 |
| Combination product | 18 | 42 | 0 | 60 |
| Total | 93 | 175 | 1 | 269 |

**Table 2.**
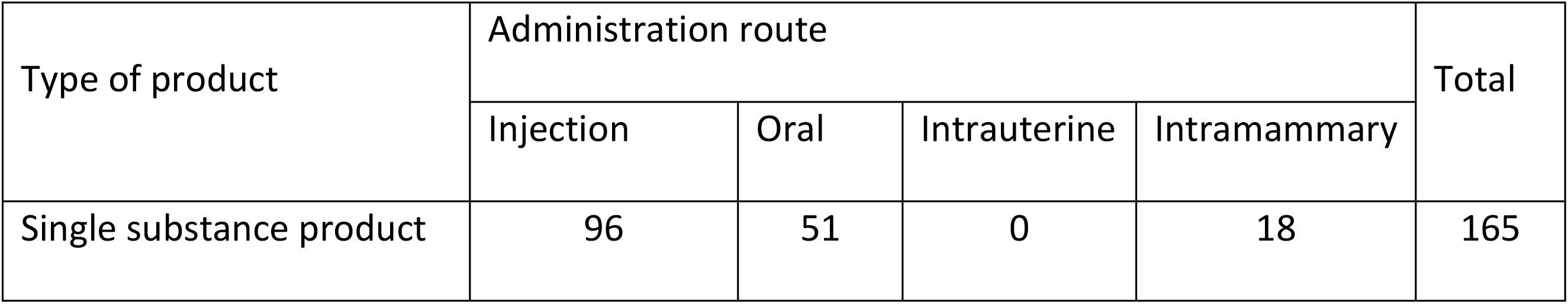
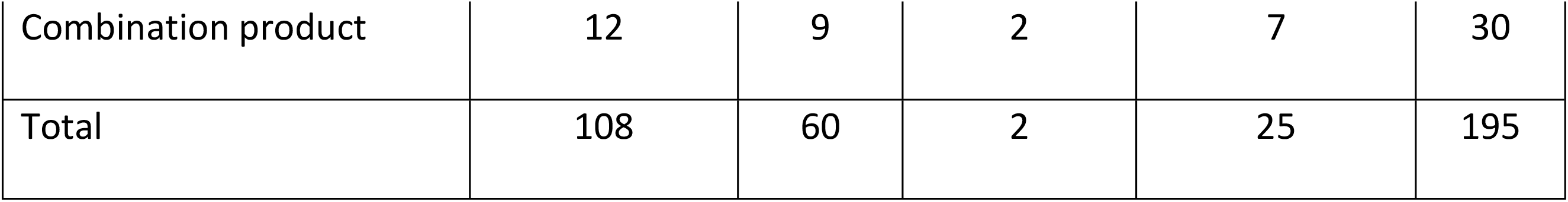
Distribution of the antimicrobial products approved for use in cattle in Japan by administration route and the number of active ingredients contained

**Table 3.**
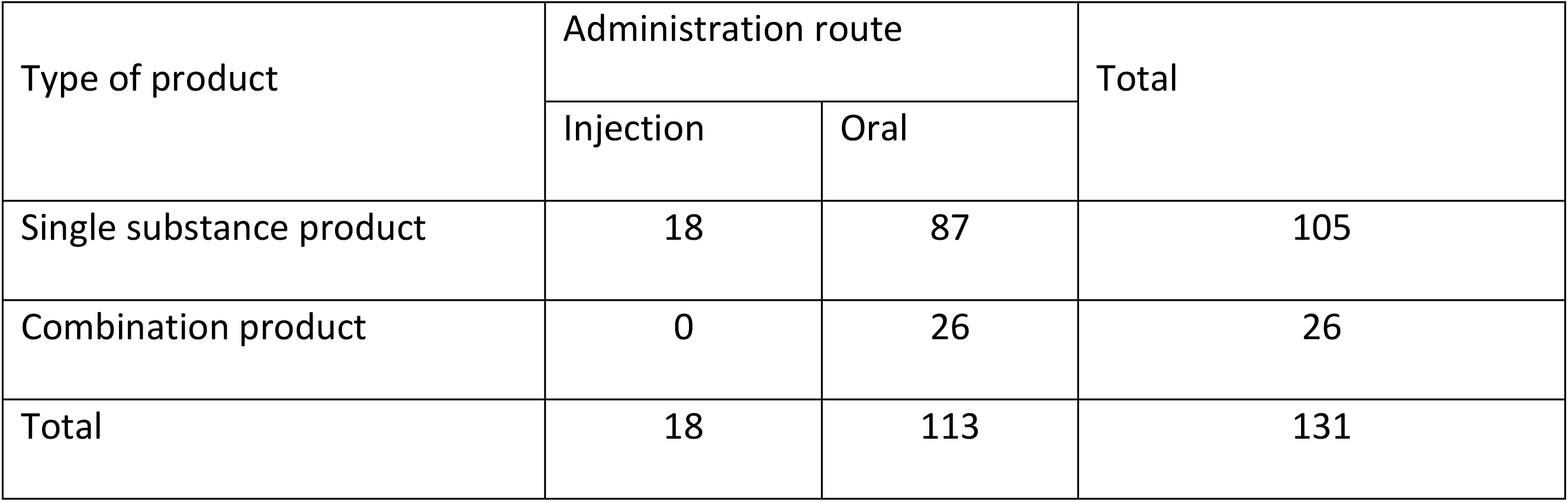
Distribution of the antimicrobial products approved for use in poultry in Japan by administration route and the number of active ingredients contained

**by administration route and the number of active ingredients contained**
| Type of product | Administration route |  | Total |
| --- | --- | --- | --- |
|  | Injection | Oral |  |
| Single substance product | 18 | 87 | 105 |
| Combination product | 0 | 26 | 26 |
| Total | 18 | 113 | 131 |

**Table 4.** Japanese DDD values (DDDjp) defined in this study for antimicrobial agents used for pigs, cattle and poulty in Japan and corresponding DDD values (DDDvet) defined by the European Medicines Agency

| Antimicrobial class | Antimicrobial agent<br>(active ingredient) | Product type | Administration<br>route | DDDvet | DDDjp | Number of<br>products |
| --- | --- | --- | --- | --- | --- | --- |
| Antimicrobial agents used for pigs |  |  |  |  |  |  |
| Tetracyclines | Oxytetracycline | Single substance product | Injection | 7.5 | 6.5 | 2 |
| Amphenicols | Thiamphenicol | Single substance product | Injection | 75.0 | 20.0 | 2 |
|  | Florfenicol | Single substance product | Injection | 9.5 | 5.0 | 4 |
| Penicillins | Ampicillin | Single substance product | Injection | 12.0 | 6.5 | 7 |
|  | Amoxicillin | Single substance product | Injection | 8.9 | 7.5 | 1 |
|  | Procaine Benzylpenicillin | Single substance product | Injection | 12.0 | 2.7 | 8 |
|  | Procaine Bensylpenicillin | Combination product | Injection | 5.4 | 7.2 | 6 |
| Cefalosporins | Ceftiofur | Single substance product | Injection | 3.0 | 2.5 | 5 |
|  | Cefquinome | Single substance product | Injection | 1.9 | 1.5 | 2 |
|  | Sulfadimethoxine | Single substance product | Injection | 30.0 | 60.0 | 2 |
| Sulfonamides | Sulfamonomethoxine | Single substance product | Injection |  | 70.0 | 1 |
|  | Sulfadoxine | Combination product | Injection | 14.0 | 30.0 | 3 |
|  | Trimethoprim | Combination product | Injection | 3.0 | 6.0 | 3 |
| Trimethoprim | Erythromycin | Single substance product | Injection | 21.0 | 4.5 | 0 |
| Macrolides | Tylosin | Single substance product | Injection | 13.0 | 6.0 | 3 |
|  | Tulathromycin | Single substance product | Injection |  | 2.5 | 4 |
| Lincosamides | Lincomycin | Single substance product | Injection | 10.0 | 7.5 | 4 |
| Aminoglycosides | Dihydrostreptomycin | Single substance product | Injection | 20.0 | 60.0 | 2 |
|  | Dihydrostreptomycin | Combination product | Injection |  | 15.0 | 6 |
|  | Kanamycin | Single substance product | Injection | 28.0 | 15.0 | 11 |
|  | Kanamycin (topical) | Single substance product | 局所 |  | 110.0 | 1 |
| Quinolones | Enrofloxacin | Single substance product | Injection | 3.4 | 2.6 | 8 |
|  | Danofloxacin | Single substance product | Injection | 1.2 | 1.3 | 1 |
|  | Marbofloxacin | Single substance product | Injection | 2.0 | 2.0 | 3 |
|  | Orbifloxacin | Single substance product | Injection |  | 3.8 | 3 |
| Pleuromutilins | Tiamulin | Single substance product | Injection | 12.0 | 10.0 | 2 |
| Tetracyclines | Doxycycline | Single substance product | Oral | 11.0 | 9.0 | 10 |
|  | Chlortetracycline | Single substance product | Oral | 31.0 | 10.8 | 6 |
|  | Chlortetracycline | Combination product | Oral |  | 6.0 | 2 |
|  | Oxytetracycline | Single substance product | Oral | 26.0 | 9.4 | 7 |
|  | Oxytetracycline | Combination product | Oral |  | 7.0 | 1 |
| Amphenicoles | Thianphenicol | Single substance product | Oral | 35.0 | 5.0 | 7 |
|  | Florfenicol | Single substance product | Oral | 10.0 | 1.5 | 18 |
| Penicillins | Ampicillin | Single substance product | Oral | 30.0 | 8.0 | 8 |
|  | Amoxicillin | Single substance product | Oral | 17.0 | 6.5 | 8 |
|  | Benzylpenicillin | Combination product | Oral |  | 0.8 | 5 |
| Sulfonamides | Sulfadimethoxine | Single substance product | Oral | 48.0 | 54.0 | 2 |
|  | Sulfadimethoxine | Combination product | Oral | 24.0 | 28.8 | 2 |
|  | Sulfamonomethoxine | Single substance product | Oral |  | 40.0 | 7 |
|  | Sulfamonomethoxine | Combination product | Oral | 9.4 | 8.1 | 4 |
|  | Sulfamethoxine | Combination product | Oral | 20.0 | 4.7 | 7 |
|  | Sulfadimidine | Combination product | Oral | 23.0 | 6.0 | 2 |
| Trimethoprim | Trimethoprim | Combination product | Oral | 4.7 | 1.4 | 9 |
|  | Ormethoprim | Combination product | Oral |  | 2.7 | 4 |
| Macrolides | Tylosin | Single substance product | Oral | 12.0 | 11.3 | 13 |
|  | Tilmicosin | Single substance product | Oral | 15.0 | 5.0 | 9 |
|  | Tylvalosin | Single substance product | Oral | 3.6 | 1.4 | 2 |
|  | Mirosamycin | Single substance product | Oral |  | 2.5 | 0 |
| Lincosamides | Lincomycin | Single substance product | Oral | 7.6 | 4.2 | 6 |
| Aminoglycosides | Streptomycin | Single substance product | Oral |  | 20.0 | 1 |
|  | Streptomycin | Combination product | Oral |  | 4.2 | 3 |
|  | Gentamicin | Single substance product | Oral | 1.4 | 0.6 | 1 |
|  | Kanamycin | Combination product | Oral |  | 4.2 | 2 |
|  | Apramycin | Single substance product | Oral | 9.0 | 4.0 | 1 |
|  | Fragiomycin | Combination product | Oral |  | 4.9 | 1 |
| Fluoroquinolones | Norfloxacin | Single substance product | Oral |  | 7.5 | 1 |
|  | Oxolinic acid | Single substance product | Oral | 26.0 | 20.0 | 3 |
| Other quinolones | Tiamulin | Single substance product | Oral | 9.7 | 6.4 | 15 |
| Pleuromutilins | Valnemulin | Single substance product | Oral | 5.3 | 2.6 | 1 |
| Polymyxins | Colistin | Single substance product | Oral | 5.0 | 4.8 | 7 |
| Total for pigs |  |  |  |  |  | 269 |
| Antimicrobial agents used for cattle |  |  |  |  |  |  |
| Tetracyclines | Oxytetracycline | Single substance product | Injection | 6.5 | 6.0 | 7 |
| Amphenicols | Thiamphenicol | Single substance product | Injection |  | 20.0 | 2 |
|  | Florfenicol | Single substance product | Injection | 13.0 | 10.0 | 9 |
| Penicillins | Ampicillin | Single substance product | Injection | 11.0 | 6.2 | 14 |
|  | Amoxicillin | Single substance product | Injection | 8.3 | 7.5 | 1 |
|  | Benzylpenicillin | Single substance product | Injection | 14.0 | 2.1 | 1 |
|  | Procaine benzylpenicillin | Single substance product | Injection | 13.0 | 7.5 | 8 |
|  | Procaine benzylpenicillin | Combination product | Injection |  | 4.2 | 6 |
|  | Aspoxicillin | Single substance product | Injection |  | 6.3 | 0 |
| Cefalosporins | Cefazolin | Single substance product | Injection |  | 5.0 | 4 |
|  | Ceftiofur | Single substance product | Injection | 1.0 | 2.3 | 5 |
|  | Cefquinome | Single substance product | Injection | 1.5 | 1.0 | 2 |
| Sulfonamides | Sulfadimethoxine | Single substance product | Injection | 30.0 | 35.0 | 2 |
|  | Sulfamonomethoxine | Single substance product | Injection |  | 25.0 | 1 |
| Macrolides | Tylosin | Single substance product | Injection | 13.0 | 7.0 | 3 |
|  | Tulathromycin | Single substance product | Injection | 0.3 | 2.5 | 3 |
|  | Tilmicosin | Single substance product | Injection | 4.0 | 10.0 | 5 |
| Aminoglycosides | Dihydrostreptomycin | Single substance product | Injection | 25.0 | 15.0 | 3 |
|  | Dihydrostreptomycin | Combination product | Injection |  | 8.8 | 6 |
|  | Kanamycin | Single substance product | Injection | 15.0 | 7.5 | 11 |
| Quinolones | Enrofloxacin | Single substance product | Injection | 4.2 | 4.2 | 8 |
|  | Danofloxacin | Single substance product | Injection |  | 1.3 | 1 |
|  | Marbofloxacin | Single substance product | Injection | 3.6 | 2.0 | 3 |
|  | Orbifloxacin | Single substance product | Injection |  | 3.8 | 3 |
| Others | Fosfomycin | Single substance product | Injection |  | 15.0 | 1 |
| Tetracyclines | Chlortetracycline | Single substance product | Oral | 22.0 | 12.5 | 6 |
|  | Oxytetracycline | Single substance product | Oral | 20.0 | 8.1 | 7 |
|  | Oxytetracycline | Combination product | Oral |  | 12.5 | 1 |
| Penicillins | Ampicillin | Single substance product | Oral | 29.0 | 8.0 | 8 |
|  | Amoxicillin | Single substance product | Oral | 20.0 | 6.5 | 8 |
| Sulfonamides | Sulfamonomethoxine | Single substance product | Oral |  | 45.0 | 8 |
|  | Sulfamonomethoxine | Combination product | Oral |  | 11.3 | 2 |
| Trimethoprim | Ormethoprim | Combination product | Oral |  | 3.8 | 2 |
| Macrolides | Tylosin | Single substance product | Oral | 41.0 | 30.8 | 1 |
|  | Tilmicosin | Single substance product | Oral | 21.0 | 14.1 | 2 |
| Aminoglycosides | Streptomycin | Single substance product | Oral | 70.0 | 20.0 | 1 |
|  | Gentamicin | Single substance product | Oral | 7.0 | 2.0 | 1 |
|  | Fragiomyacin | Combination product | Oral | 7.4 | 8.8 | 1 |
| Fluoroquinolones | Enrofloxacin | Single substance product | Oral | 4.7 | 3.8 | 1 |
| Other quinolones | Oxolinic acid | Single substance product | Oral | 17.0 | 15.0 | 4 |
| Polymyxins | Colistin | Single substance product | Oral | 4.8 | 3.5 | 2 |
| Others | Fosfomycin | Single substance product | Oral |  | 30.0 | 2 |
| Penicillins | Procaine benzylpenicillin | Combination product | Intrauterine |  | 0.2 | 1 |
| Aminoglycosides | Dihydrostreptomycin | Combination product | Intrauterine |  | 0.3 | 1 |
| Cephalosporins | Cefazolin | Single substance product | Intramammary |  | 0.3 | 9 |
|  | Cefalonium | Single substance product | Intramammary |  | 0.2 | 3 |
|  | Cefuroxime | Single substance product | Intramammary |  | 0.6 | 2 |
| Tetracyclines | Oxytetracycline | Single substance product | Intramammary |  | 1.1 | 1 |
| Penicillins | Procaine benzylpenicillin | Combination product | Intramammary |  | 0.4 | 4 |
|  | Dicloxacillin | Single substance product | Intramammary |  | 0.3 | 2 |
| Aminoglycosides | Dihydrostreptomycin | Combination product | Intramammary |  | 0.6 | 2 |
|  | Kanamycin | Combination product | Intramammary |  | 0.5 | 1 |
|  | Fradiomycin | Combination product | Intramammary |  | 0.5 | 1 |
| Lincosamides | Pirlimycin | Single substance product | Intramammary |  | 0.1 | 1 |
| Total for cattle |  |  |  |  |  | 196 |
| Antimicrobial agents used for poultry |  |  |  |  |  |  |
| Tetracyclines | Oxytetracycline | Single substance product | Injection |  | 31.3 | 5 |
| Aminoglycosides | Dihydrostreptomycin | Single substance product | Injection |  | 62.5 | 2 |
|  | Kanamycin | Single substance product | Injection |  | 37.5 | 11 |
| Tetracyclines | Doxycycline | Single substance product | Oral | 15.0 | 17.3 | 12 |
|  | Chlortetracycline | Single substance product | Oral | 30.0 | 35.1 | 6 |
|  | Oxytetracycline | Single substance product | Oral | 39.0 | 40.7 | 7 |
| Amphenicoles | Thianphenicol | Single substance product | Oral | 55.0 | 39.0 | 7 |
|  | Florfenicol | Single substance product | Oral | 30.0 | 20.0 | 1 |
| Penicillins | Amoxicillin | Single substance product | Oral | 16.0 | 30.0 | 8 |
|  | Ampicillin | Single substance product | Oral | 108.0 | 16.3 | 8 |
|  | Procaine benzylpenicillin | Combination product | Oral |  | 4.7 | 5 |
| Sulfonamides | Sulfadimethoxine | Single substance product | Oral | 65.0 | 106.3 | 2 |
|  | Sulfadimethoxine | Combination product | Oral | 31.0 | 56.2 | 2 |
|  | Sulfamonomethoxine | Single substance product | Oral |  | 97.5 | 7 |
|  | Sulfamonomethoxine | Combination product | Oral | 9.5 | 31.9 | 4 |
|  | Sulfamethoxazole | Combination product | Oral | 27.0 | 32.5 | 2 |
| Trimethoprim | Trimethoprim | Combination product | Oral | 6.4 | 6.4 | 4 |
|  | Ormethoprim | Combination product | Oral |  | 10.5 | 4 |
| Macrolides | Tylosin | Single substance product | Oral | 81.0 | 78.9 | 13 |
|  | Tylvalosin | Single substance product | Oral | 25.0 | 47.6 | 3 |
| Lincosamides | Lincomycin | Single substance product | Oral | 8.6 | 5.1 | 6 |
| Aminoglycosides | Streptomycin | Combination product | Oral |  | 23.4 | 3 |
|  | Kanamycin | Combination product | Oral |  | 23.4 | 2 |
| Fluoroquinolones | Norfloxacin | Single substance product | Oral |  | 20.0 | 1 |
|  | Enrofloxacin | Single substance product | Oral | 10.0 | 11.5 | 1 |
|  | Ofloxacin | Single substance product | Oral |  | 7.5 | 1 |
| Other quinolones | Oxolinic acid | Single substance product | Oral | 20.0 | 39.7 | 4 |
| Total for poultry |  |  |  |  |  | 131 |

DDDvet: DDD values in mg/kg/day defined by the European Medicines Agency (EMA)
DDDjp: DDD values in mg/kg/day defined in this study using DDD values of antimicrobial products approved and marketed for use in Japan

### Comparison of the DDDjp values with DDDvet values for antimicrobial agens for use in pigs

A comparison of 44 pairs of DDDjp and DDDvet values of antimicrobial agents for use in pigs was possible. The distribution of the logarithmic quotients of daily doses is shown in Figure 1A. A total of 27 compared values showed deviations of more than 50% (Fig. 1A). A significant difference between the DDDvet and DDDjp values was observed for antimicrobial agents used for pigs (P <0.01) with the median QDDD value of 0.66.

**Fig 1.**
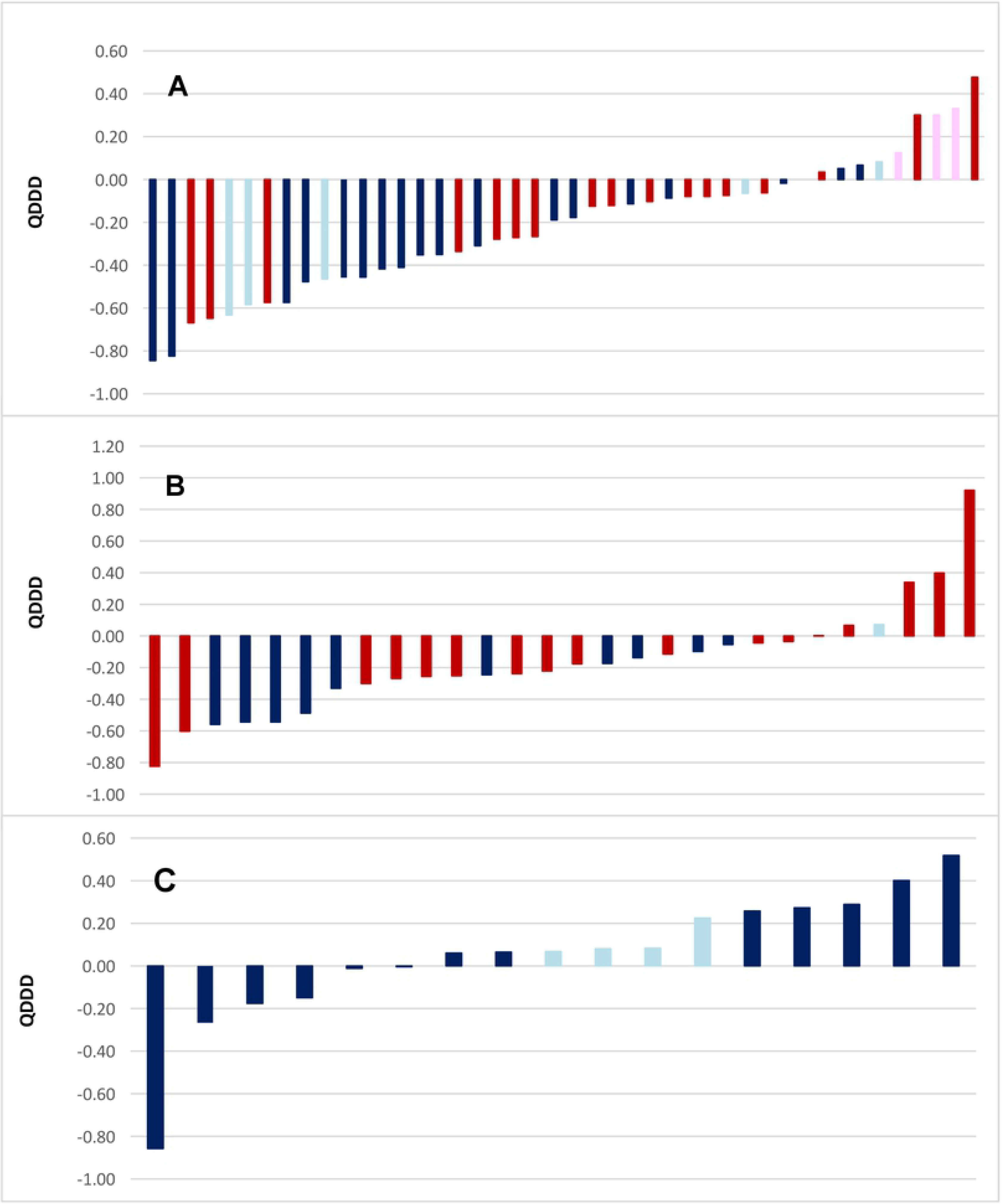
Comparison of the DDDjp values of antimicrobial agents approved in Japan with the corresponding values of the European Medicines Agency. Bars indicate an antimicrobial agent approved for use in pigs (A), cattle (B) and poultry (C) for which both DDDjp and DDDvet values are available with QDDD values in ascending order. The red, pink, blue and light blue bars are injection/single substance, injection/combination, oral/single and oral/combination respectively.

The administration route showed a significant difference (P = 0.011) in terms of the level of the deviations between the DDDjp and DDDvet values (Table 5). In comparison with the guideline values of the EMA, the injection solutions with the active substances kanamycin and ampicillin showed lower daily doses with QDDD of 0.54 and 0.54 respectively. Injection solution with the active substance benzyl-penicillin showed lower daily doses (QDDD=0.23) in single substance products but showed higher daily doses (QDDD=1.33) in combination products. With antimicrobial agents for oral administration, florfenicol, trimethoprim and tiamulin were approved in Japan with lower daily doses with QDDD of 0.15, 0.31 and 0.66 respectively (Table 6).

**Table 5.**
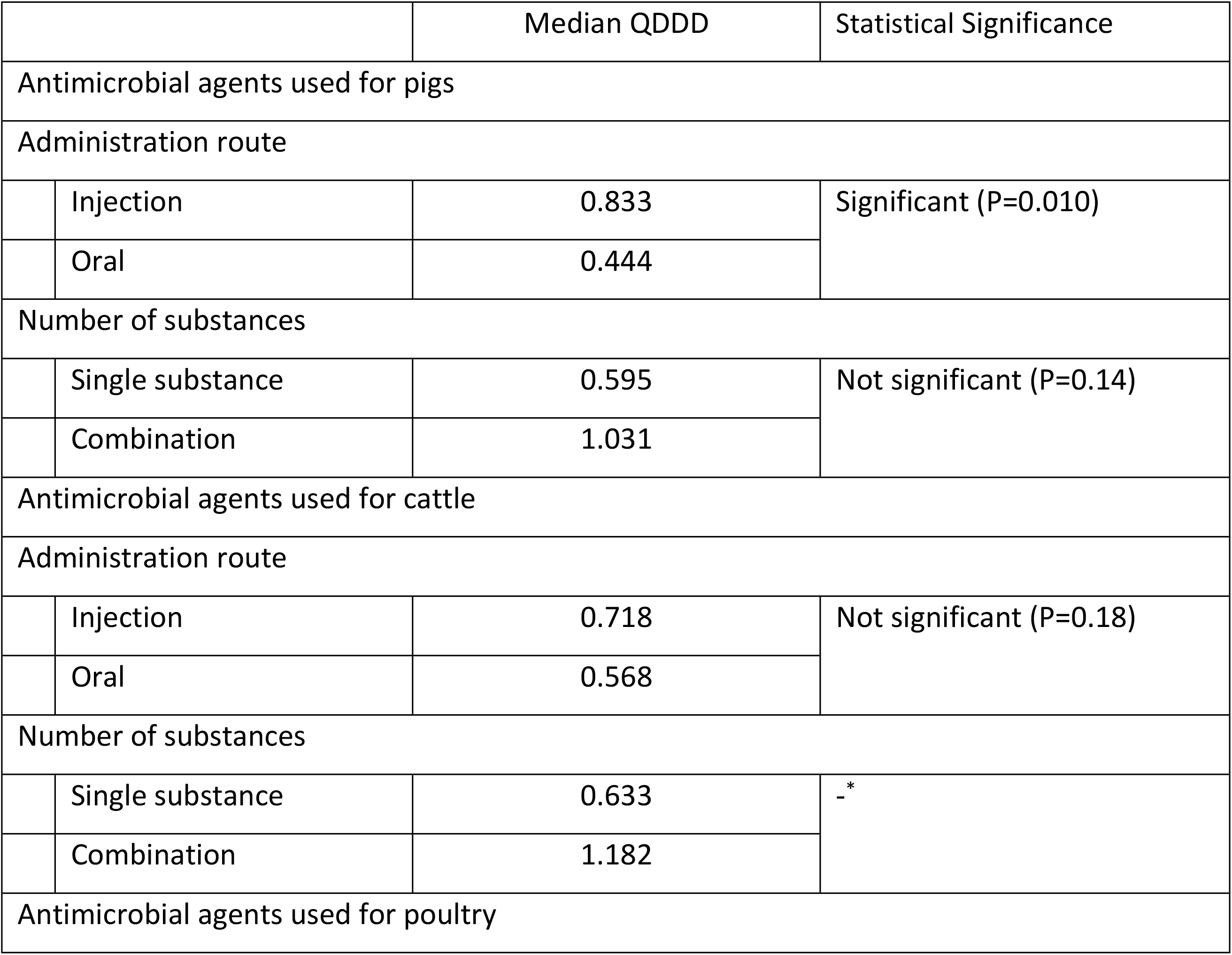
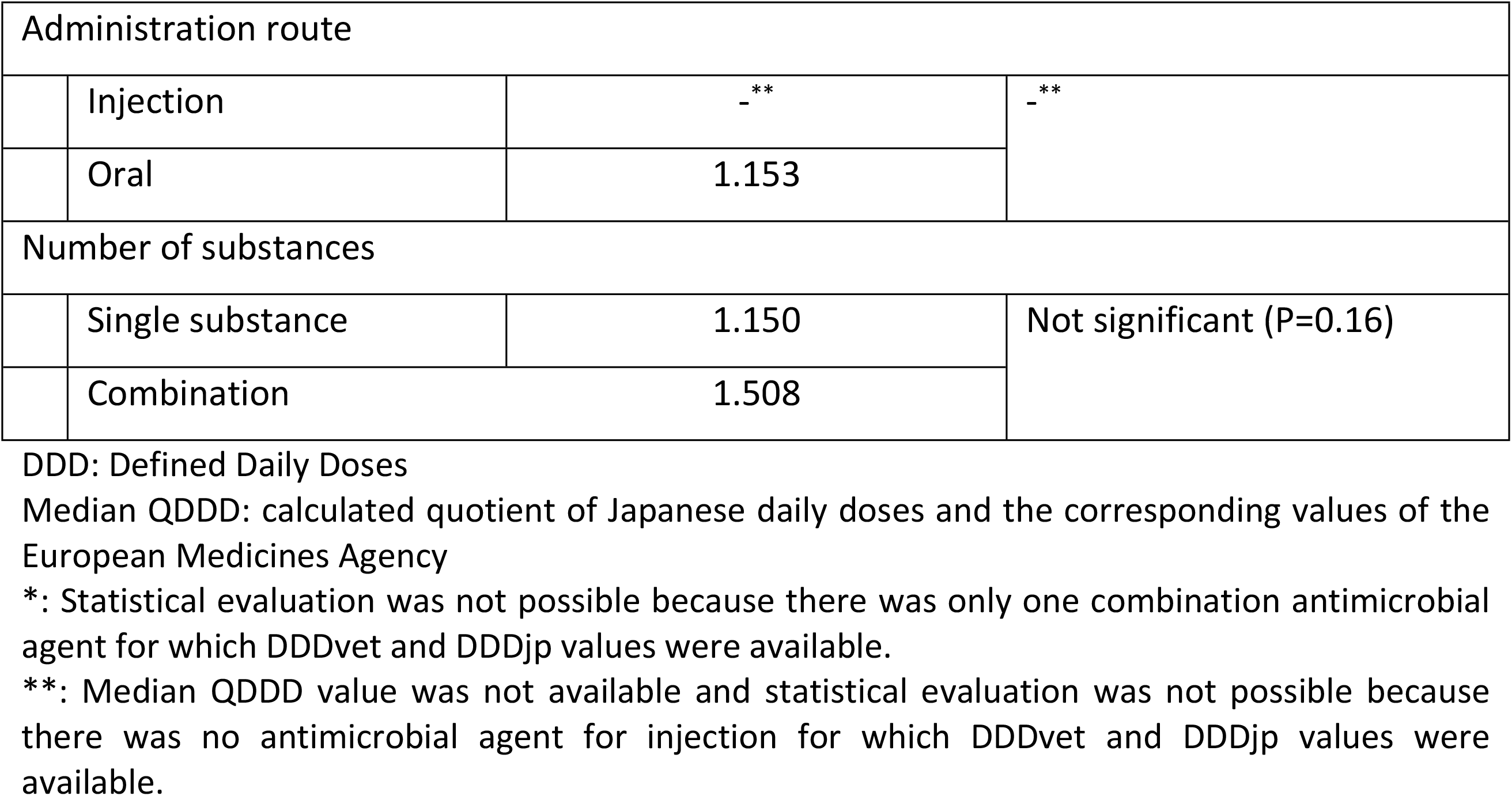
Statistical evaluation of the calculated quotients QDDD in relation to administration routes and the number of active ingredients contained in the preparation (Mann-Whitney U Test)

|  |  | Median QDDD | Statistical Significance |
| --- | --- | --- | --- |
| Antimicrobial agents used for pigs |  |  |  |
| Administration route |  |  |  |
|  | Injection | 0.833 | Significant (P=0.010) |
|  | Oral | 0.444 |  |
| Number of substances |  |  |  |
|  | Single substance | 0.595 | Not significant (P=0.14) |
|  | Combination | 1.031 |  |
| Antimicrobial agents used for cattle |  |  |  |
| Administration route |  |  |  |
|  | Injection | 0.718 | Not significant (P=0.18) |
|  | Oral | 0.568 |  |
| Number of substances |  |  |  |
|  | Single substance | 0.633 | - * |
|  | Combination | 1.182 |  |
| Antimicrobial agents used for poultry |  |  |  |

| Administration route |  |  |  |
| --- | --- | --- | --- |
|  | Injection | - ** | - ** |
|  | Oral | 1.153 |  |
| Number of substances |  |  |  |
|  | Single substance | 1.150 | Not significant (P=0.16) |
|  | Combination | 1.508 |  |

DDD: Defined Daily Doses
Median QDDD: calculated quotient of Japanese daily doses and the corresponding values of the European Medicines Agency
\*: Statistical evaluation was not possible because there was only one combination antimicrobial agent for which DDDvet and DDDjp values were available.
\*\*: Median QDDD value was not available and statistical evaluation was not possible because there was no antimicrobial agent for injection for which DDDvet and DDDjp values were available.

**Table 6.** Average daily doses (DDDjp) of the most frequently approved antimicrobial agents (divided by administration routes) in the pig, cattle and poultry sector in Japan and their comparison with the values of the European Medicines Agency (DDDvet) based on calculated quotients (QDDD)

| Administration route | Antimicrobial agent (active ingredient) | Number of substances | DDDjp (Number of products approved) | DDDvet | QDDD |
| --- | --- | --- | --- | --- | --- |
| Antimicrobial agents used for pigs |  |  |  |  |  |
| Injection | Kanamycin | single substance | 15.0 (11) | 28.0 | 0.54 |
|  | Procaine benzylpenicillin | single substance | 2.7 (8) | 12.0 | 0.23 |
|  | Enrofloxacin | single substance | 2.6 (8) | 3.4 | 0.76 |
|  | Ampicillin | single substance | 6.5 (7) | 12.0 | 0.54 |
|  | Procaine benzylpenicillin | combination | 7.2 (6) | 5.4 | 1.33 |
| Oral | Florfenicol | single substance | 1.5 (18) | 10.0 | 0.15 |
|  | Tiamulin | single substance | 6.4 (15) | 9.7 | 0.66 |
|  | Tylosin | single substance | 14.0 (13) | 12.0 | 1.17 |
|  | Doxycycline | single substance | 9.0 (10) | 11.0 | 0.82 |
|  | Trimethoprim | combination | 1.4 (9) | 4.7 | 0.31 |
| Antimicrobial agents used for cattle |  |  |  |  |  |
| Injection | Ampicillin | single substance | 6.2 (14) | 11.0 | 0.56 |
|  | Kanamycin | single substance | 7.5 (11) | 15.0 | 0.50 |
|  | Florfenicol | single substance | 10.0 (9) | 13.0 | 0.77 |
|  | Procaine benzylpenicillin | single substance | 7.5 (8) | 13.0 | 0.58 |
|  | Enrofloxacin | single substance | 4.2 (8) | 4.2 | 1.00 |
| Oral | Ampicillin | single substance | 8.0 (8) | 29.0 | 0.28 |
|  | Amoxicillin | single substance | 6.5 (8) | 20.0 | 0.33 |
|  | Oxytetracycline | single substance | 8.1 (7) | 20.0 | 0.41 |
|  | Chlortetracycline | single substance | 12.5 (6) | 22.0 | 0.57 |
|  | Oxolinic acid | single substance | 15.0 (4) | 17.0 | 0.88 |
| Antimicrobial agents used for poultry |  |  |  |  |  |
| Oral | Tylosin | single substance | 78.9 (13) | 81.0 | 0.97 |
|  | Doxycycline | single substance | 17.3 (12) | 15.0 | 1.15 |
|  | Ampicillin | single substance | 16.3 (8) | 108.0 | 0.15 |
|  | Amoxicillin | single substance | 30.0 (8) | 16.0 | 1.88 |
|  | Oxytetracycline | single substance | 40.7 (7) | 39.0 | 1.04 |

DDDjp: defined daily doses of the active ingredient used in pigs in Japan established in this study. DDDvet: defined daily doses of the active ingredient used in pig in Europe established by the European Medicines Agency. QDDD: calculated quotient of DDDjp/DDDvet.

The number of active substances contained in a veterinary medicinal product (P <0.01) did not show a significant difference in terms of the level of the deviations between the daily doses DDDjp and DDDvet (Table 5).

### Comparison of the DDDjp values with DDDvet values for antimicrobial agents for use in cattle

A comparison of 27 pairs of DDDjp and DDDvet values of antimicrobial agents for use in cattle was possible. The distribution of the logarithmic quotients of daily doses is shown in Figure 1B. A total of 10 compared values showed deviations of more than 50% (Fig. 1B). A significant difference between the DDDvet and DDDjp values was observed for antimicrobial agents used for cattle (P <0.01) with the median QDDD value of 0.63.

Neither the administration route nor the number of active substance contained in a veterinary medicinal product showed a significant difference in terms of the level of the deviations between the DDDjp and DDDvet values (Table 5). In comparison with the guideline values of the EMA, the injection solutions with the active substances ampicillin, kanamycin and procaine bensylpenicillin showed lower daily doses with QDDD of 0.56, 0.50 and 0.58 respectively. With antimicrobial agents for oral administration, ampicillin, amoxlillin and oxytetracycline were approved in Japan with lower daily doses with QDDD of 0.28, 0.33 and 0.41 respectively (Table 6).

### Comparison of the DDDjp values with DDDvet values for antimicrobial agents for use in poultry

A comparison of 17 pairs of DDDjp and DDDvet values of antimicrobial agents for use in poultry was possible. The distribution of the logarithmic quotients of daily and treatment dosages is shown in Figure 1C. A total of 5 compared values showed deviations of more than 50% (Fig. 1C). No statistically significant difference was observed between the DDDvet and DDDjp values with the median QDDD value of 1.17.

Neither the administration route nor the number of active substances contained in a veterinary medicinal product showed a significant difference in terms of the level of the deviations between the DDDjp and DDDvet values (Table 5). In comparison with the guideline values of the EMA, the oral solutions with the active substance ampicillin showed lower daily dose with QDDD of 0.15 and amoxicillin with higher daily dose with QDDD of 1.88 (Table 6).

## Discussion

The present study defined national daily dosages (DDDjp) for the first time for all antimicrobial agents used in products approved for use in pigs, cattle and poultry in Japan. A comparison with corresponding values of the EMA was possible for most antimicrobial agents. The comparison within this study shows that the medians of DDDjp and DDDvet values differ significantly, and that DDDjp values of some antimicrobial have considerable deviations from corresponding DDDvet values. Canada also found that in developing their country-specific DDD values, the majority of their DDD values were lower than their corresponding DDDvet values [14]. In a previous study conducted by Echtermann defining Swiss daily doses (DDDch), the difference between DDDch and DDDvet values was not as significant as between DDDjp and DDDvet values in the current study [15]. There are many reasons for the difference observed between DDDvet and DDDjp or DDD values in other non-European countries. One reason is that the EMA might have had a wider range of antimicrobial doses to work with due to the collection of antimicrobial agent doses from nine European countries [9, 13]. The different labelling regulations, different treatment indications and different husbandry practices might all contribute to the variations in DDDvet and DDDjp values. However, fully elucidating the reasons for these differences is beyond the scope of this study.

Furthermore, this study showed that DDDvets did not cover all the antimicrobial agents used in veterinary medicine in Japan. Although drawing conclusions from differences between assigned DDDjp and DDDvet values is difficult, the difference between DDDvet and DDDjp values and absence of DDDvet values for some antimicrobial agents marketed in Japan indicate that DDDjp rather than DDDvet should be used as the basis for the calculation of antimicrobial use monitoring in farm animals in Japan, assuming that DDDjp better reflects the actual dosage used in food-producing animals in Japan. This is a reasonable assumption considering that Japanese veterinarians are more likely to follow dosage instructions rather than European instructions when they treat pigs using antimicrobials marketed in Japan.

To determine if the use of DDDjp is really recommendable as the basis for the calculation of antimicrobial use on farms in Japan, application of DDDjp and DDDvet values for calculation of the numbers of DDDs for comparison using actual antimicrobial usage data on farms is indispensable, but this will be a subject of future studies.

## Acknowledgements

This study was conducted as part of the research project on Regulatory research projects for food safety, animal health and plant protection (Grant number: JPJ008617.17935699. 1120) funded by the Ministry of Agriculture, Forestry and Fisheries of Japan.

